# Associations of age, sex, and bicuspid aortic valve with biaxial mechanical properties and circumferential residual stress in ascending thoracic aortic aneurysm

**DOI:** 10.64898/2026.10.01.755974

**Authors:** Noor Ghadie, Munir Boodhwani, Alexis Mccreath Frangakis, Navneet Kang, Aryam Toubar, Michel Labrosse

## Abstract

**Background:** Ascending thoracic aortic aneurysm (ATAA) is characterized by progressive biomechanical remodeling that is incompletely captured by maximum aortic diameter. The relationships between age, sex, aortic valve phenotype (BAV vs TAV), and aneurysmal mechanics remains incompletely understood.

**Methods:** Ascending aortic specimens from 54 patients undergoing elective ATAA repair were evaluated. Biaxial testing quantified Green strain and local stiffness at membrane tensions of 60 and 120 N/m. Opening angle was used to assess circumferential residual stress and stretch. Associations with age were evaluated using Pearson correlations, and differences between sex and valve phenotype were also explored. Multivariable linear regression models including age, sex, and valve phenotype were used to evaluate their adjusted associations with each mechanical outcome.

**Results:** Increasing age was associated with increased local stiffness and reduced Green strain and remained the strongest determinant of biaxial mechanical properties after adjustment. Compared with tricuspid aortic valve specimens, BAV specimens exhibited lower Green strain together with lower local stiffness. Females demonstrated greater Green strain in unadjusted analyses; however, after adjustment, sex remained associated only with longitudinal Green strain at 60 N/m. None of the evaluated clinical characteristics consistently explained opening angle or residual stretch.

**Conclusions:** Age was the predominant clinical determinant of biaxial mechanical remodeling in ATAA, whereas valve phenotype demonstrated selective associations and sex contributed comparatively little after adjustment. Circumferential residual stress appeared largely independent of these clinical characteristics, suggesting that residual stress and biaxial mechanics capture complementary aspects of aneurysmal remodeling.

## 1 Introduction

Ascending thoracic aortic aneurysm (ATAA), a disease with increasing prevalence over the years, often progresses to catastrophic events such as aortic dissection or rupture^1–3^. Although maximum aortic diameter remains the principal criterion for recommending prophylactic surgery^4^, it incompletely captures the mechanical and structural properties of the aortic wall and does not fully explain patient-specific differences in disease progression. Given the central role of aortic wall mechanics in governing aortic failure, there is a critical need to better characterize the mechanical properties of aneurysmal tissue to improve our understanding of disease progression and ultimately refine patient risk stratification^5–8^.

The mechanical properties of the ascending aorta depend on both its nonlinear load-bearing response and its residual stress state. During the cardiac cycle, the characteristics of the nonlinear stress–strain behavior of the aorta dictate its ability to expand during systole and recoil during diastole, contributing to the Windkessel effect by storing and subsequently releasing elastic energy to maintain continuous blood flow^9^. In parallel, circumferential residual stress is thought to redistribute in-vivo wall stresses through the vessel thickness under physiological loading, thereby reducing stress concentrations within the wall^10,11^. As aneurysms develop, both the nonlinear biaxial stress-strain response of the tissue and its residual stress state undergo progressive remodeling^12^. Understanding the factors that contribute to these mechanical changes is therefore important for improving our understanding of aneurysmal remodeling.

Patient-specific clinical characteristics are thought to contribute to the mechanical heterogeneity observed in patients with ATAA. Among these, age, sex, and valve phenotype have received particular attention. Aging is accompanied by progressive extracellular matrix remodeling, including elastin degradation, collagen accumulation, and increased collagen cross-linking, all of which are expected to influence the mechanical behavior of the aortic wall^13^. Bicuspid aortic valve (BAV), the most common congenital heart disease, affects approximately 1–2% of the population, and nearly half of affected individuals develop ATAA^14^. Additionally, although ATAA is more common in men, women experience poorer clinical outcomes, including higher risks of dissection and rupture and acute aortic syndromes at smaller aneurysm diameters even after correcting for body size^5,15^.

Although previous studies have suggested that age, sex, and valve phenotype may influence the mechanical properties of the aorta during ATAA, the available evidence remains limited and, in some cases, conflicting^6–8,16^. The objective of this study was to evaluate the associations of age, sex, and valve phenotype with biaxial mechanical properties and circumferential residual stress in patients with ATAA.

## 2 Methods

### 2.1 Study Population and Tissue Preparation

The analytic cohort consisted of 54 patients and encompassed age and aneurysm-size ranges that minimized potential confounding between sex and valve phenotype groups. Baseline characteristics are summarized in Table 1. Written informed consent was obtained from all participants, and the study was approved by the University of Ottawa Heart Institute Research Ethics Board (protocol 20140762-01H, approved December 22, 2014).

**Table 1.** Baseline characteristics of the analytic cohort.

| Characteristic | Full Cohort<br>(n=54) | Male<br>(n=36) | Female<br>(n=18) | p-value | TAV<br>(n=33) | BAV<br>(n=20) | p-value |
| --- | --- | --- | --- | --- | --- | --- | --- |
| Age (years) | 60.0 ± 12.5 | 61.5 ± 9.7 | 57.2 ± 16.6 | 0.327 | 60.8 ± 12.6 | 58.8 ± 12.4 | 0.583 |
| Male, n (%) | 36 (66.7%) | — | — | — | 19 (57.6%) | 16 (80.0%) | 0.137 |
| BAV, n (%) | 20 (37.0%) | 16 (44.4%) | 4 (22.2%) | 0.137 | — | — | — |
| AAD (mm) | 46.0 ± 9.0 | 46.6 ± 6.2 | 45.0 ± 13.0 | 0.622 | 46.1 ± 10.3 | 46.0 ± 6.8 | 0.982 |
| Weight (kg) | 90.9 ± 25.6 | 98.9 ± 27.5 | 77.7 ± 15.1 | **0.001** | 90.9 ± 22.4 | 91.1 ± 31.9 | 0.981 |
| Height (cm) | 174.8 ± 9.2 | 176.6 ± 6.2 | 171.8 ± 12.4 | 0.144 | 174.6 ± 9.9 | 175.3 ± 8.0 | 0.785 |
| BSA (m<sup>2</sup>) | 2.10 ± 0.31 | 2.19 ± 0.29 | 1.92 ± 0.25 | **<0.001** | 2.11 ± 0.29 | 2.09 ± 0.34 | 0.835 |
| Smoking, n (%) | 27 (50.0%) | 19 (52.8%) | 8 (44.4%) | 0.240 | 17 (51.5%) | 10 (50.0%) | 0.758 |
| Hypertension, n (%) | 27 (50.0%) | 18 (50.0%) | 9 (50.0%) | 0.558 | 21 (63.6%) | 6 (30.0%) | 0.122 |

Values are mean ± standard deviation or n (%). Continuous variables were compared using Welch's t-test; categorical variables were compared using Fisher's exact test. Percentages are based on available data; therefore, denominators vary between variables due to missing values. BAV = bicuspid aortic valve; AAD = maximum ascending aortic diameter; BSA = body surface area.

Fresh tubular shaped ascending aortic specimens were collected from patients undergoing elective ATAA repair. Following surgical excision, specimens were either tested immediately or stored at 4°C and tested within 24 hours. Owing to differences in tissue availability and specimen suitability for each experimental protocol, the number of specimens included in each of the mechanical tests varied and is reported throughout the manuscript.

### 2.2 Biaxial Mechanical Testing

Up to four 1 x 1 cm square samples were extracted along the circumference of 43 ATAA patients. The samples were mounted to a 5N capacity CellScale biaxial machine (BioTester 5000, CellScale, Waterloo, Canada) using four tungsten rakes, each with 5 tines. Specimens were subjected to 10 preconditioning testing cycles followed by nine biaxial testing protocols as described previously^17^. In this work, only the equibiaxial tension testing protocol was considered for each case in the post-processing phase explained next.

The image tracking module in LabJoy was used to process the images and data for tension-strain plots. Nodal coordinates in the reference undeformed configuration (*X*_1_, *X*_2_, *X*_3_), and current deformed configuration, (*x*_1_, *x*_2_, *x*_3_), were used to determine the components of the deformation gradient

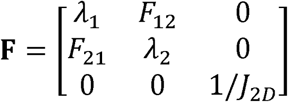

where *λ*_1_ and *λ*_2_ are the in-plane stretch ratios, *F*_12_ and *F*_21_ are shear deformation components, and *J*_2*D*_ = *λ*_1_*λ*_2_ - *F*_12_*F*_21_ (incompressibility assumption). The Green–Lagrange strain tensor was subsequently determined as **E = ½(F^T^F−I)**, with **I** being the identity tensor. Using the measured forces *f*_1_ and *f*_2_ and undeformed dimensions between the tines of the CellScale tester *L*_1_ and *L*_2_, in the circumferential and longitudinal directions respectively, second Piola-Kirchhoff (PK) membrane tensions were computed as

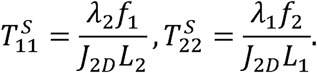

Second PK membrane tension–Green strain curves were then built, and the mechanical features were extracted at predefined equibiaxial membrane tensions of 60 and 120 N/m. Stiffness in each loading direction was calculated as the local slope of the second PK membrane tension versus Green strain relationship at the specified reference tensions.

### 2.3 Opening Angle

One to three aortic rings of about 3-4 mm in height were extracted from the distal region of 50 tubular ATAA segments. Rings were immersed in isotonic saline (300 mOsm NaCl) over a 1 × 1 cm calibration grid and imaged in the unloaded (closed) configuration. Each ring was then radially cut, allowing the release of circumferential residual stress and opening toward its nearly stress-free configuration. After a 30-minute equilibration period, the opened rings were re-imaged.

Opening angle measurements were obtained from the paired images using a custom MATLAB program described previously^11^. Following image calibration, the inner and outer contours were manually traced in both the closed and opened configurations. Assuming that the midline arc length as well as the thickness are the same in the unstressed and unloaded configurations, the opening angle was computed as 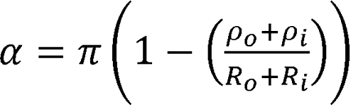, where *ρ*_i_ and *ρ*_o_ are the inner and outer radii in the closed configurations and *R*_i_ and *R*_o_ in the opened stress-free configurations. When the ring opens beyond the flat configuration, the open radii are assigned negative values, allowing opening angles greater than 180° to be represented. For specimens with multiple rings, opening angle was averaged to obtain a single representative value for each patient.

### 2.4 Residual Stretch

Treating the aortic wall as one homogeneous layer, the residual stretch in the circumferential direction can be calculated as 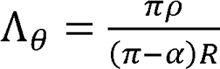 ^18,19^, where - *ρ* and *R* are the radii in the closed and open configurations respectively such that *ρ*_i_ < *ρ* < *ρ*_o_ and *R*_i_ < *R* < *R*_o_. The inner residual stretch was calculated as the circumferential residual stretch at the intima. The residual stretch difference between the innermost and outermost layers was calculated as the difference between the residual stretch at the adventitia and the residual stretch at the intima.

### 2.5 Statistical Analysis

Statistical testing was performed using Python 3.14.4. Associations between age and mechanical properties were evaluated using Pearson correlation coefficients. For comparisons between sex and valve phenotype groups, normality was assessed using the Shapiro–Wilk test and visual inspection of Q–Q plots. Normally distributed variables were compared using Welch’s t-test, whereas non-normally distributed variables were compared using the Mann–Whitney U test.

Multivariable linear regression models were used to evaluate the independent associations of age, sex, and valve phenotype with each mechanical property. Age was modeled as a continuous variable, whereas sex (male = 0, female = 1) and valve phenotype (TAV = 0, BAV = 1) were included as binary predictors. Regression coefficients (β), p-values, and model R^2^ are reported. The unique contribution of each predictor was quantified using partial R², calculated by comparing the full model with a reduced model excluding the predictor of interest.

All tests were two-sided, and p < 0.05 was considered statistically significant.

## 3 Results

### 3.1 Associations with Age

Association between age and mechanical properties assessed via Pearson’s correlation analysis are summarized in Figure 1 and Figure 2. Increasing age was associated with greater circumferential stiffness at both 60 N/m (r = 0.722, p < 0.001) and 120 N/m (r = 0.512, p < 0.001), as well as greater longitudinal stiffness at 60 N/m (r = 0.358, p = 0.020) and 120 N/m (r = 0.435, p = 0.004). Similarly, increasing age was associated with lower circumferential strain at 60 N/m (r = −0.446, p = 0.003) and 120 N/m (r = −0.670, p < 0.001), and lower longitudinal strain at 120 N/m (r = −0.467, p = 0.002). Longitudinal strain at 60 N/m demonstrated a similar near-significant trend (r = −0.275, p = 0.078). In contrast, as shown in Figure 2, age was not significantly associated with opening angle, inner residual stretch, or residual stretch difference (all p ≥ 0.396).

**Figure 1.**
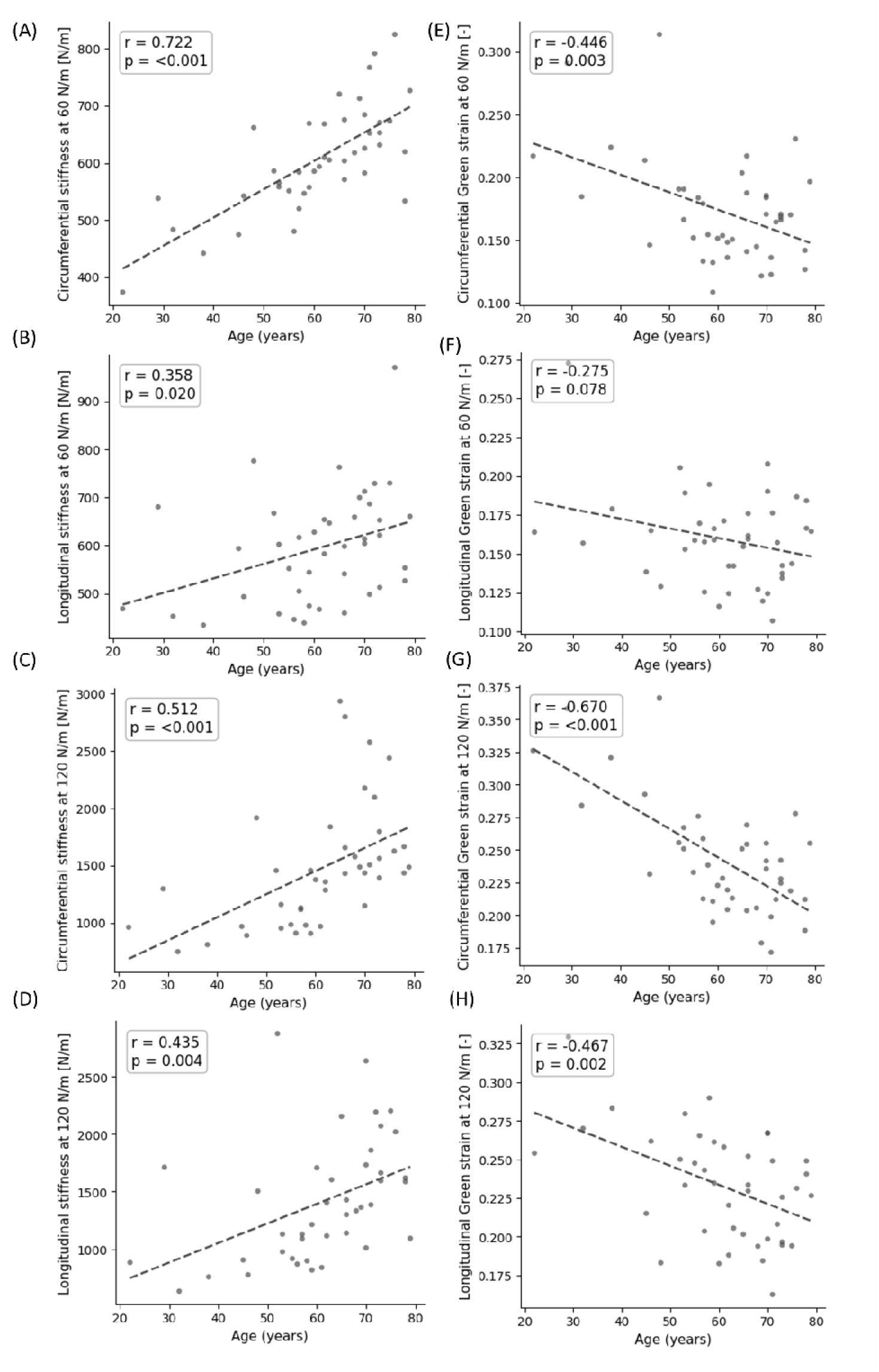
Pearson correlations between age and circumferential stiffness at (A) 60 N/m and (B) 120 N/m, longitudinal stiffness at (C) 60 N/m and (D) 120 N/m, circumferential Green strain at (E) 60 N/m and (F) 120 N/m, longitudinal Green strain at (G) 60 N/m and (H) 120 N/m (n = 42).

**Figure 2.**
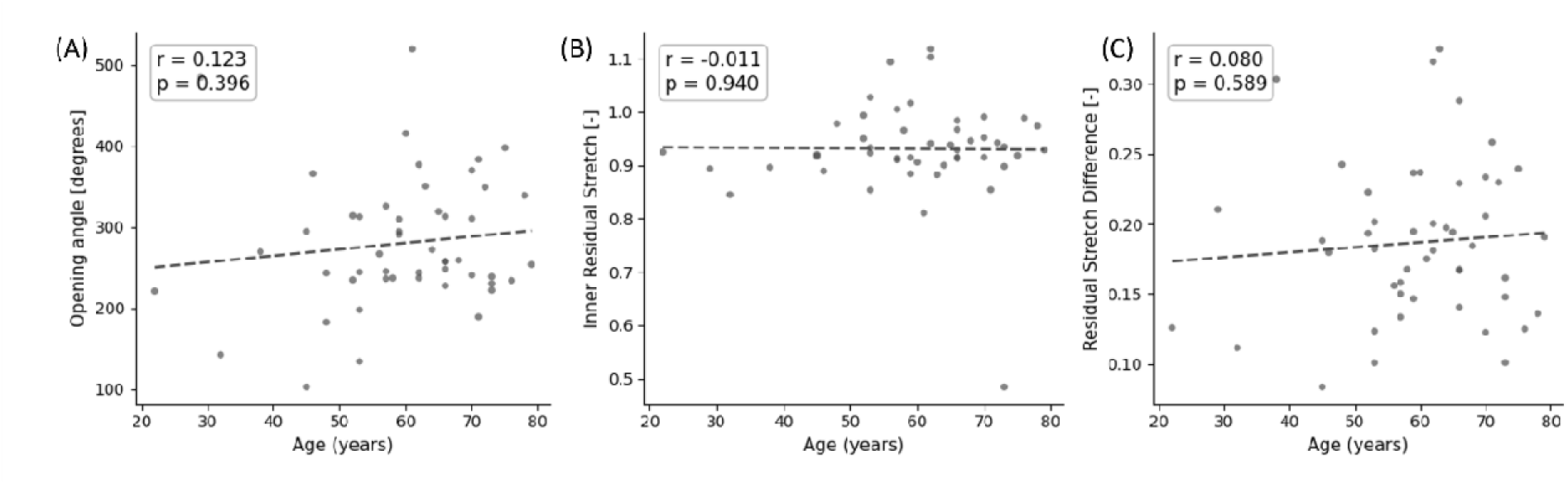
Pearson correlations between age and (A) opening angle (n = 50), (B) inner residual stretch (n = 48), (C) residual stretch difference (n = 48).

### 3.2 Differences between Sexes

No significant sex-related differences in tangent stiffness were observed at either loading level (Figure 3A,B). At 60 N/m, circumferential stiffness was 621 ± 108 kPa in males and 611 ± 124 kPa in females (p = 0.969), while longitudinal stiffness was 601 ± 129 kPa and 612 ± 138 kPa, respectively (p = 0.806). At 120 N/m, circumferential stiffness was 1492 ± 569 kPa in males and 1521 ± 561 kPa in females (p = 0.766), while longitudinal stiffness was 1376 ± 446 kPa and 1536 ± 690 kPa, respectively (p = 0.437).

**Figure 3.**
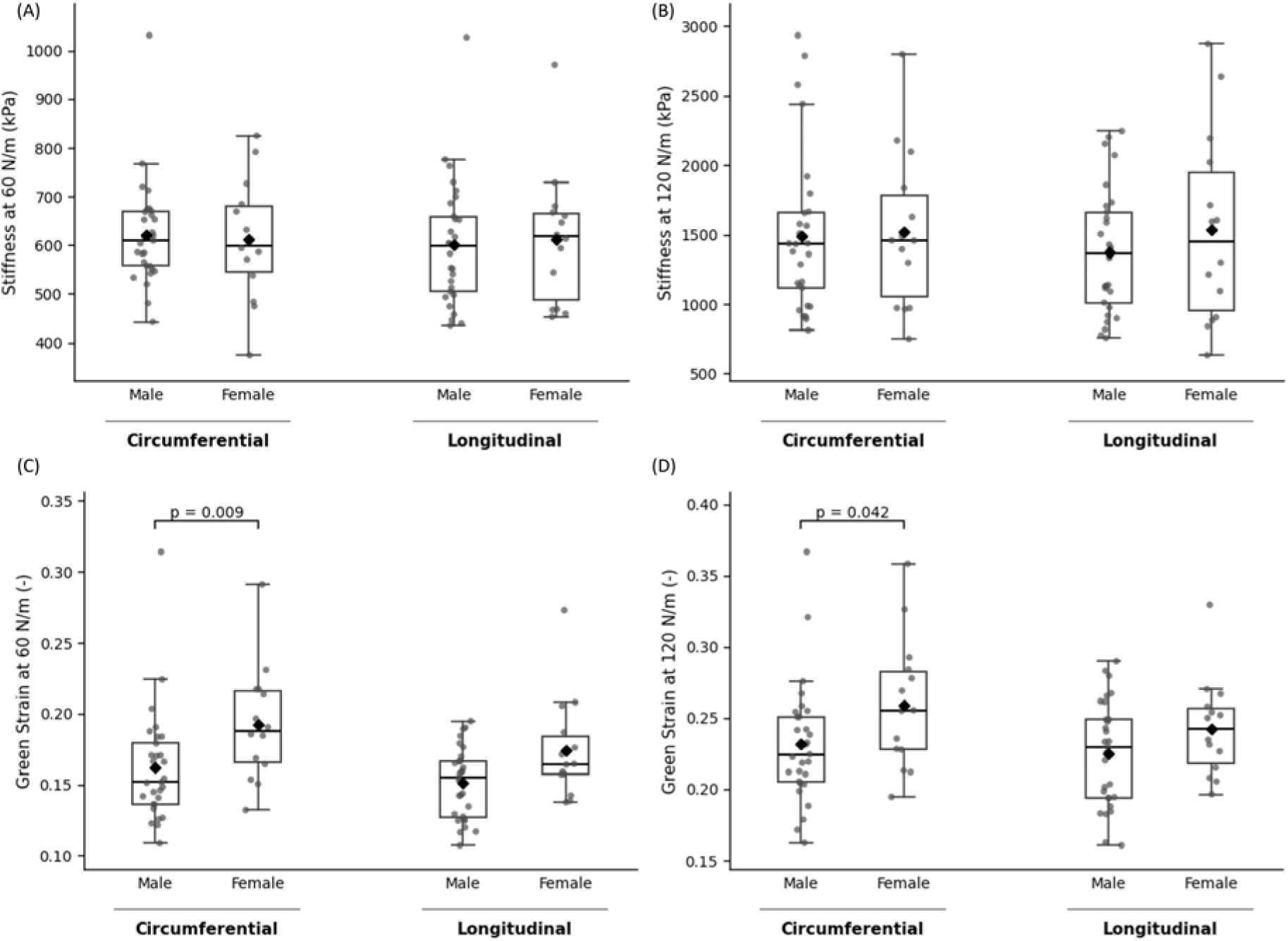
Comparisons of biaxial tensile properties according to sex: male (n = 29) vs. female (n = 14). Stiffness at (A) 60 N/m, and (B) 120 N/m, Green strain at (C) 60 N/m, and (D) 120 N/m. Boxplots represent the median (center line), interquartile range (box), and 1.5 × interquartile range (whiskers). Black diamonds indicate the group mean, and gray points represent individual specimens. p-values are displayed for statistically significant results only.

In contrast, females exhibited greater circumferential Green strain than males at both 60 N/m (0.193 ± 0.040 vs. 0.162 ± 0.040, p = 0.009; Figure 3C) and 120 N/m (0.259 ± 0.046 vs. 0.232 ± 0.042, p = 0.042; Figure 3D). Longitudinal Green strain was numerically greater in females than males at 60 N/m (0.174 ± 0.036 vs. 0.151 ± 0.025, p = 0.064; Figure 3C) and 120 N/m (0.243 ± 0.034 vs. 0.225 ± 0.037, p = 0.134; Figure 3D), but neither difference was statistically significant.

Opening angle was similar between males and females (280 ± 68° vs. 282 ± 101°, p = 0.941; Figure 4A). Similarly, no significant sex-related differences were observed in inner residual stretch (0.952 ± 0.063 vs. 0.919 ± 0.052, p = 0.073; Figure 4B) or residual stretch difference (0.191 ± 0.054 vs. 0.181 ± 0.055, p = 0.540; Figure 4C).

**Figure 4.**
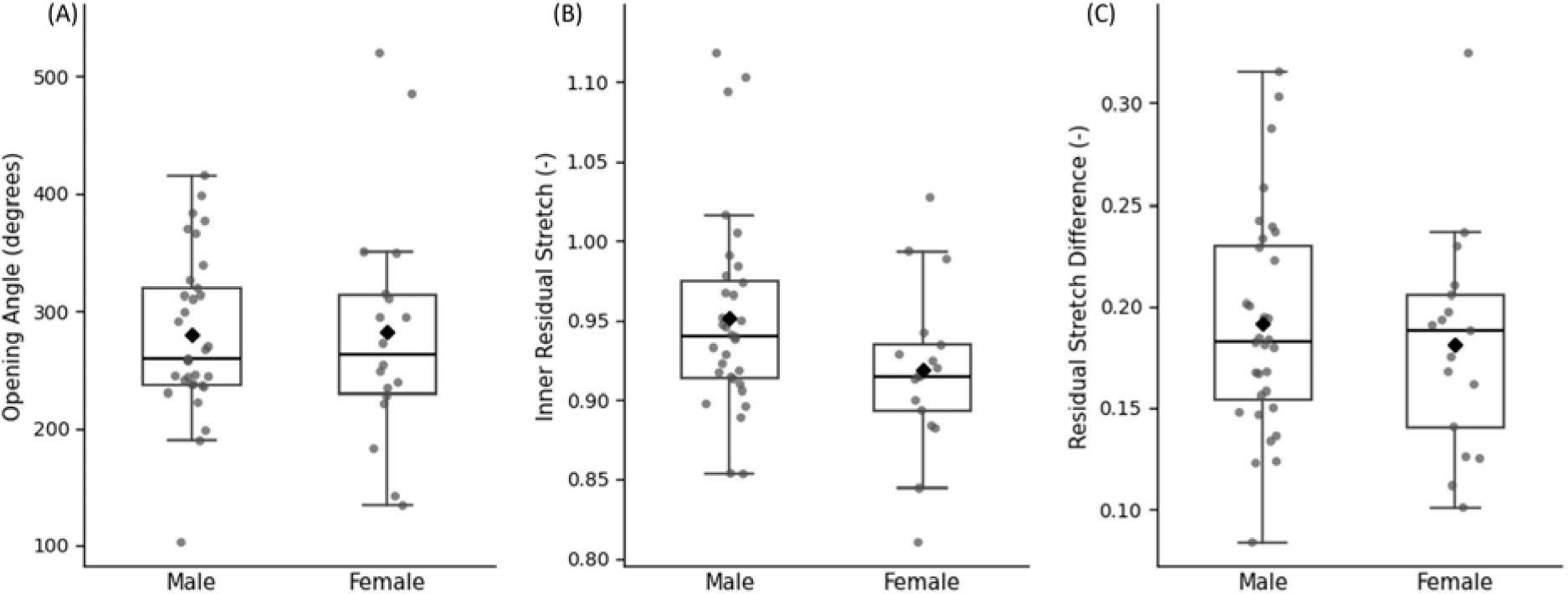
Comparisons of residual stress and stretch according to sex (male vs. female). (A) Opening angle (male, n = 33; female, n = 18), (B) inner residual stretch (male, n = 32; female, n = 17), (C) residual stretch difference (male, n = 32; female, n = 17). Boxplots represent the median (center line), interquartile range (box), and 1.5 × interquartile range (whiskers). Black diamonds indicate the group mean, and gray points represent individual specimens. p-values are displayed for statistically significant results only.

### 3.3 Differences between Valve Phenotypes

Circumferential stiffness did not differ significantly between BAV and TAV specimens at 60 N/m (595 ± 92 vs. 616 ± 94 kPa, p = 0.463; Figure 5A), but was lower in BAV at 120 N/m (1240 ± 383 vs. 1613 ± 561 kPa, p = 0.018; Figure 5B). Longitudinal stiffness was lower in BAV than TAV specimens at both 60 N/m (545 ± 92 vs. 625 ± 117 kPa, p = 0.022; Figure 5A) and 120 N/m (1191 ± 413 vs. 1542 ± 549 kPa, p = 0.024; Figure 5B).

**Figure 5.**
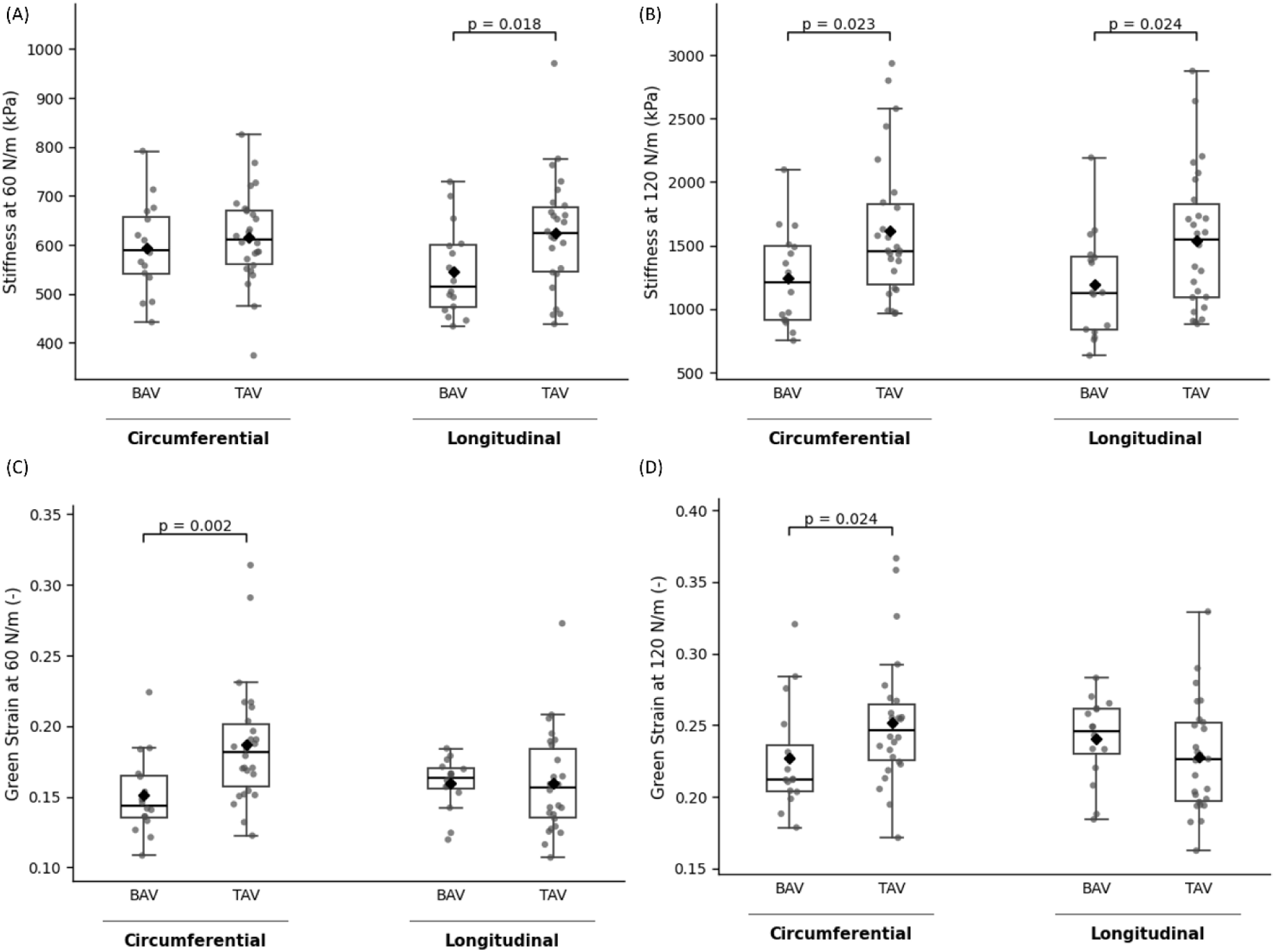
Comparisons of biaxial tensile properties according to valve phenotype: BAV (n = 16) vs. TAV (n = 26). Stiffness at (A) 60 N/m, and (B) 120 N/m, Green strain at (C) 60 N/m and (D) 120 N/m. Boxplots represent the median (center line), interquartile range (box), and 1.5 × interquartile range (whiskers). Black diamonds indicate the group mean, and gray points represent individual specimens. p-values are displayed for statistically significant results only.

Circumferential Green strain was lower in BAV than TAV specimens at both 60 N/m (0.151 ± 0.029 vs. 0.187 ± 0.044, p = 0.002; Figure 5C) and 120 N/m (0.227 ± 0.038 vs. 0.252 ± 0.045, p = 0.024; Figure 5D). In contrast, longitudinal Green strain did not differ significantly between BAV and TAV specimens at either 60 N/m (0.159 ± 0.018 vs. 0.160 ± 0.036, p = 0.623; Figure 5C) or 120 N/m (0.241 ± 0.028 vs. 0.227 ± 0.039, p = 0.218; Figure 5D).

No significant differences in circumferential residual stress or stretch were observed according to valve phenotype (Figure 6). Opening angle was 290 ± 96° in BAV and 275 ± 72° in TAV specimens (p = 0.337; Figure 6A). Similarly, inner residual stretch was 0.937 ± 0.082 in BAV and 0.942 ± 0.047 in TAV specimens (p = 0.678; Figure 6B), while residual stretch difference was 0.177 ± 0.056 and 0.195 ± 0.053, respectively (p = 0.286; Figure 6C).

**Figure 6.**
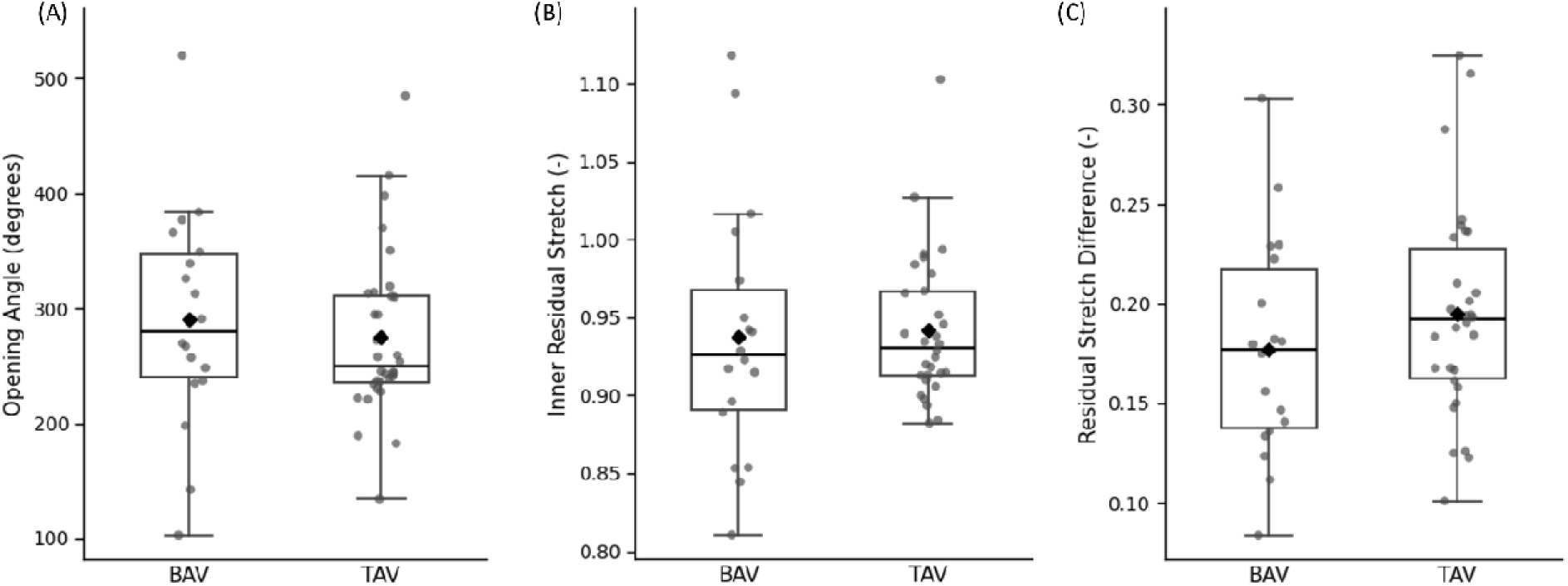
Comparisons of residual stress and stretch according to valve phenotype. (A) Opening angle (BAV, n = 18; TAV, n = 32), (B) inner residual stretch (BAV, n = 18; TAV, n = 30), (C) residual stretch difference (BAV, n = 18; TAV, n = 30). Boxplots represent the median (center line), interquartile range (box), and 1.5 × interquartile range (whiskers). Black diamonds indicate the group mean, and gray points represent individual specimens. p-values are displayed for statistically significant results only.

### 3.4 Multivariable Associations

To evaluate the independent associations of age, sex, and valve phenotype with mechanical remodeling, multivariable regression analyses were performed (Table 2). Overall, the multivariate analyses were consistent with the univariate findings, with age demonstrating the most consistent independent association across the evaluated mechanical properties.

**Table 2.** Regression model parameters.

| Mechanical Feature | Age $\beta$ | p | BAV $\beta$ | p | Sex $\beta$ | p | $R^2$ |
| --- | --- | --- | --- | --- | --- | --- | --- |
| **Circumferential stiffness at 60 N/m** | 5.184 | **<0.001** | -7.093 | 0.739 | 32.18 | 0.156 | 0.553 |
| **Longitudinal stiffness at 60 N/m** | 3.091 | **0.016** | -69.89 | **0.045** | 26.93 | 0.452 | 0.248 |
| **Circumferential stiffness at 120 N/m** | 20.33 | **<0.001** | -318.4 | **0.032** | 111.5 | 0.463 | 0.374 |
| **Longitudinal stiffness at 120 N/m** | 18.11 | **0.002** | -273.3 | 0.073 | 226.1 | 0.154 | 0.321 |
| **Circumferential Green strain at 60 N/m** | -0.001363 | **0.002** | -0.0346 | **0.003** | 0.01284 | 0.273 | 0.413 |
| **Longitudinal Green strain at 60 N/m** | -0.0004665 | 0.181 | 0.003921 | 0.681 | 0.02038 | **0.047** | 0.168 |
| **Circumferential Green strain at 120 N/m** | -0.002187 | **<0.001** | -0.02683 | **0.011** | 0.005991 | 0.572 | 0.554 |
| **Longitudinal Green strain at 120 N/m** | -0.001113 | **0.005** | 0.01448 | 0.170 | 0.01266 | 0.252 | 0.269 |
| **Opening angle** | 0.976 | 0.317 | 21.43 | 0.399 | 10.8 | 0.670 | 0.031 |
| **Inner residual stretch** | 0.000547 | 0.454 | -0.0098 | 0.606 | -0.03402 | 0.080 | 0.090 |
| **Residual stretch difference** | 0.000332 | 0.613 | -0.01943 | 0.259 | -0.01416 | 0.412 | 0.049 |

*Regression coefficients are reported in the native units of each outcome. Age coefficients represent the change per additional year. Sex was coded as male=0 and female=1, and valve phenotype as TAV=0 and BAV=1. $R^2$ represents the proportion of outcome variance explained by the complete model.*

Increasing age was independently associated with greater circumferential and longitudinal stiffness at both 60 and 120 N/m (all p ≤ 0.016), as well as lower circumferential Green strain at 60 and 120 N/m and lower longitudinal Green strain at 120 N/m (all p ≤ 0.005). In contrast, age was not associated with longitudinal Green strain at 60 N/m (p = 0.181) or with opening angle, inner residual stretch, or residual stretch difference (all p ≥ 0.317).

Valve phenotype demonstrated more selective independent associations. BAV was independently associated with lower longitudinal stiffness at 60 N/m (p = 0.045), lower circumferential stiffness at 120 N/m (p = 0.032), and lower circumferential Green strain at both 60 and 120 N/m (p = 0.003 and p = 0.011, respectively) but was not independently associated with the remaining mechanical outcomes. Sex demonstrated limited independent associations, remaining significant only for greater longitudinal Green strain at 60 N/m (p = 0.047).

Partial R² analysis further demonstrated that age consistently accounted for the largest proportion of uniquely explained mechanical variability across mechanical outcomes (Table 3). Valve phenotype contributed more modestly to selected stiffness and strain measures, whereas sex explained relatively little unique variance. All three clinical characteristics contributed minimally to the variability in opening angle and residual stretch measures.

**Table 3.**
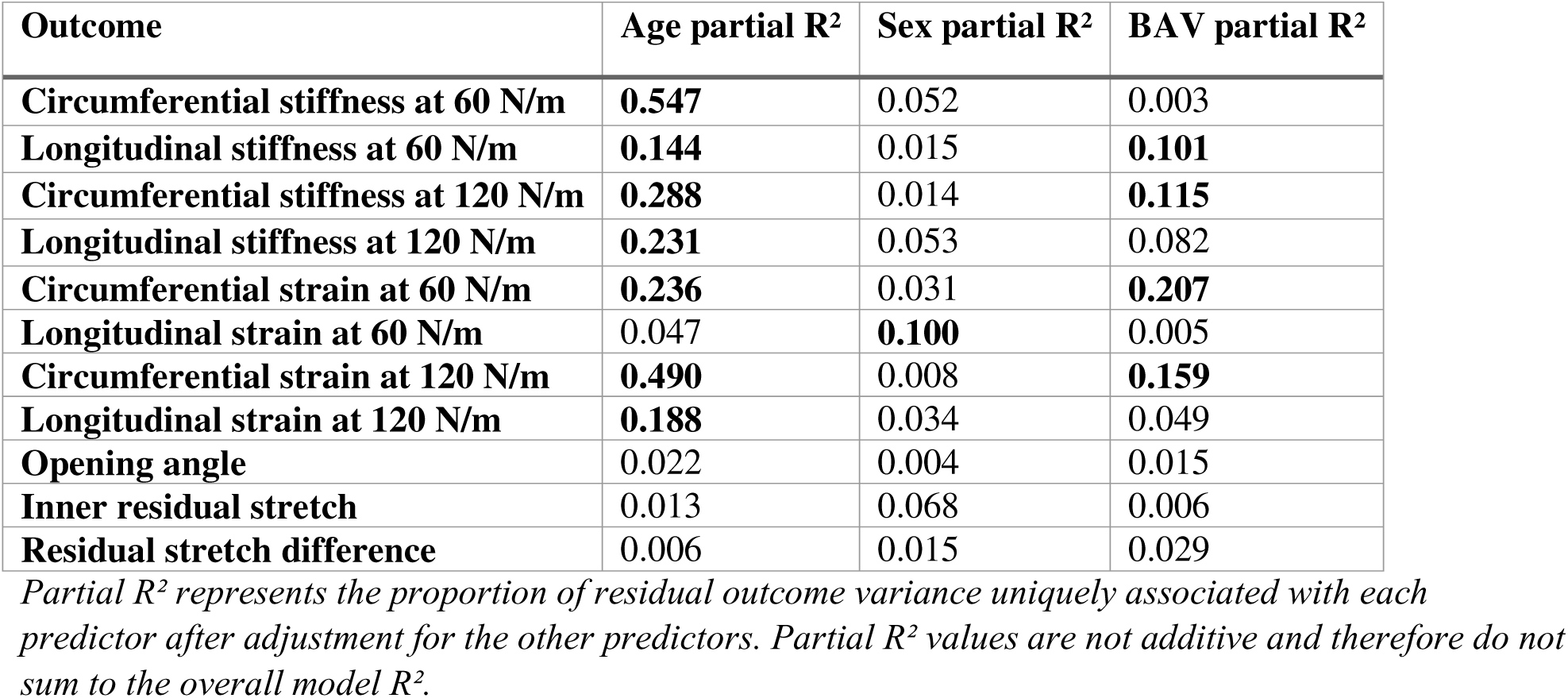
Partial R².

| Outcome | Age partial R<sup>2</sup> | Sex partial R<sup>2</sup> | BAV partial R<sup>2</sup> |
| --- | --- | --- | --- |
| **Circumferential stiffness at 60 N/m** | **0.547** | 0.052 | 0.003 |
| **Longitudinal stiffness at 60 N/m** | **0.144** | 0.015 | **0.101** |
| **Circumferential stiffness at 120 N/m** | **0.288** | 0.014 | **0.115** |
| **Longitudinal stiffness at 120 N/m** | **0.231** | 0.053 | 0.082 |
| **Circumferential strain at 60 N/m** | **0.236** | 0.031 | **0.207** |
| **Longitudinal strain at 60 N/m** | 0.047 | **0.100** | 0.005 |
| **Circumferential strain at 120 N/m** | **0.490** | 0.008 | **0.159** |
| **Longitudinal strain at 120 N/m** | **0.188** | 0.034 | 0.049 |
| **Opening angle** | 0.022 | 0.004 | 0.015 |
| **Inner residual stretch** | 0.013 | 0.068 | 0.006 |
| **Residual stretch difference** | 0.006 | 0.015 | 0.029 |

*Partial R<sup>2</sup> represents the proportion of residual outcome variance uniquely associated with each predictor after adjustment for the other predictors. Partial R<sup>2</sup> values are not additive and therefore do not sum to the overall model R<sup>2</sup>.*

## 4 Discussion

The present study evaluated the associations of age, sex, and valve phenotype with biaxial mechanical properties and circumferential residual stress in ATAA. Three principal findings emerged. First, age was the strongest and most consistent determinant of biaxial mechanical remodeling. Second, valve phenotype demonstrated more selective associations with tissue stiffness and deformation, whereas sex exhibited comparatively modest effects. Third, none of these routinely available clinical characteristics consistently explained circumferential residual stress or stretch. As such, the findings in this work indicate that different components of aneurysmal mechanics are influenced differently by patient characteristics.

### 4.1 AGE

Age was consistently associated with reduced strain and increased stiffness. Age also remained an independent predictor of strain and stiffness in multivariable analyses. The reductions in strain and increase in local stiffness provide complementary evidence of age-related mechanical stiffening of aneurysmal tissue. Consistent with these observation, age-related increases in aneurysmal tissue stiffness have been reported in ex-vivo studies^20^ and in-vivo imaging studies^21^.

In contrast to the clear age-related changes in biaxial mechanical properties, neither opening angle nor residual stretch was significantly associated with age. Moreover, an exploratory assessment of a quadratic age relationship (results not shown) did not improve the model fit, providing no evidence that opening angle followed a consistent nonlinear trajectory with age. Previous studies of dilated ascending aortas by Okamoto et al. and Sokolis et al. have reported increasing opening angle with age^22,23^. The absence of a comparable association in the present study may reflect important methodological differences between studies. Okamoto et al. determined opening angle from the two free ends and the midpoint of the inner contour, while excluding rings for which a representative angle could not be defined because much of the specimen did not remain upright or its curvature was markedly nonuniform. In contrast, the present study employed a previously validated contour-based image-processing method that traces the complete inner and outer boundaries of the opened ring before estimating opening angle from an equivalent circular representation. Although this approach retains the assumptions of circular geometry, it defines opening angle from the measured contour lengths rather than a three-point construction, allowing a broader range of opening configurations to be analyzed. Furthermore, the correlation analysis reported by Sokolis et al. appears to have incorporated measurements from multiple rings obtained from individual patients to characterize regional residual strains, whereas the present study represented each patient by a single average value. Consequently, differences in opening-angle definition, specimen selection, and the level at which observations were analyzed may have contributed to the differing age relationships.

### 4.2 Valve Phenotype

Patients with BAV exhibited lower local stiffness as well as lower Green strain compared with patients with TAV. Although lower Green strain and lower local stiffness may at first appear inconsistent, these metrics characterize different aspects of the nonlinear mechanical response. Green strain reflects the cumulative deformation achieved up to a prescribed loading state, whereas the local stiffness represents the instantaneous local slope of the curve at that operating point. Consequently, the lower strain observed in BAV indicates reduced deformation (or compliance) over the loading interval, whereas the lower stiffness indicates a more compliant local response at the specific operating point.

Previous studies support the notion that the apparent effect of valve phenotype depends on how stiffness is defined. Choudhury et al.^24^ reported a higher circumferential low-strain modulus in BAV than TAV, with no differences in the high-strain modulus, suggesting that valve-related differences were confined to the early portion of the stress-strain response. Chung et al.^6^ observed a greater tangent modulus in BAV when stiffness was evaluated at a fixed Green strain of 10%, thereby comparing tissues at an equivalent level of deformation. In contrast, Lin et al.^8^ found no significant difference between valve phenotypes using the maximum Young’s modulus, a measure of the peak stiffness attained during loading. Unlike these previous definitions, the present study quantified a local tangent stiffness at fixed physiological membrane tensions, thereby comparing tissues under equivalent loading conditions. Conclusions regarding the influence of valve phenotype on aneurysmal stiffness are therefore influenced by the mechanical definition of stiffness and the loading conditions under which it is evaluated. Together, these findings suggest that valve phenotype does not uniformly alter aneurysmal stiffness; rather, its apparent influence depends on both the mechanical metric used and the loading conditions under which stiffness is evaluated.

Valve phenotype was not associated with circumferential residual stress and stretch in the present cohort. Opening angle, inner residual stretch, and residual stretch difference did not differ between BAV- and TAV-associated aneurysms, and valve phenotype was not an independent predictor of any residual stress metric after adjustment for age and sex. These findings are consistent with previous studies reporting no significant differences in opening angle or residual strain between valve phenotypes^22,23^.

### 4.3 SEX

Unlike valve phenotype, sex had relatively modest associations with aneurysmal mechanics. Females exhibited greater deformation in unadjusted analyses, but after adjustment for age and valve phenotype only longitudinal Green strain at 60 N/m remained independently associated with sex. Previous ex-vivo studies examining sex-related differences in aneurysmal tissue mechanics have reported conflicting findings. Sokolis and Iliopoulos^16^ suggested increased wall stiffness in female compared with males. Similarly, Lin et al. found approximately 30% greater maximum Young’s modulus in females^8^. In contrast, Ferrara et al. observed no differences in maximum elastic modulus between sexes^7^. These mixed findings are mirrored by in-vivo investigations. Boczar et al. showed that carotid-femoral pulse wave velocity (a measure of aortic stiffness) did not differ between women and men with thoracic aortic aneurysms^5^, although they demonstrate that greater aortic stiffness was independently associated with faster aneurysm growth in women but not in men. Furthermore, cardiovascular magnetic resonance studies have demonstrated that the age-related increase in aortic stiffness is steeper in women than in men^25^, indicating that sex differences in aortic mechanics may be strongly influenced by age and hormonal status. Differences among studies may also reflect the populations being examined. Ex-vivo studies generally represent larger aneurysms as they evaluate patients undergoing surgical resection. Moreover, apparent sex differences may be influenced by differences in age or other clinical characteristics between groups, particularly when these factors are not accounted for in multivariable analyses. Collectively, these findings suggest that the influence of sex on aneurysmal mechanics is likely multifactorial and may depend on age, disease stage, and the mechanical metric being evaluated.

Sex did not influence opening angle, inner residual stretch, or residual stretch difference in the present cohort. In the healthy aorta, it has been reported that opening angle is higher in males compared to females^26^. In ATAA, Sokolis et al. reported larger opening angles in males than females among patients with TAV, whereas no sex-related differences were observed in BAV-associated aneurysms ^23^. In our cohort, no significant sex differences were identified within either the TAV or BAV subgroups (results not shown). The reason for this discrepancy is unclear but may reflect differences in cohort size and characteristics, study design, or statistical power.

### 4.4 Limitations

Several limitations need to be stated. Mechanical characterization was performed ex-vivo under controlled laboratory conditions. Although this approach enables detailed assessment of intrinsic tissue mechanics, it cannot fully reproduce the complex physiological environment of the living aorta. Additionally, only selected routinely available clinical characteristics were evaluated. Other factors such as genetic background, smoking, history of hypertension and inflammatory status may also contribute to patient-specific mechanical remodeling and likely account for part of the unexplained variability observed in the present study.

### 4.5 Overall implications and Conclusion

Taken together, these findings indicate that routinely available clinical characteristics explain only part of the mechanical heterogeneity observed in ATAA. Age emerged as the predominant determinant of biaxial mechanical behavior, whereas valve phenotype demonstrated more selective associations and sex contributed relatively little after adjustment. This interpretation is further supported by the partial R² analysis, which consistently demonstrated that age accounted for substantially more unique mechanical variability than either sex or valve phenotype. Age may therefore be an important consideration when interpreting aortic mechanical behavior and support further investigation into whether age-related differences in tissue mechanics contribute to susceptibility to aortic failure.

Interestingly, none of the evaluated clinical characteristics consistently explained circumferential residual stress. The lack of association between age, sex, and valve phenotype and circumferential residual stress suggests that the mechanisms governing residual stress remodeling may differ from those responsible for changes in biaxial stiffness and deformation. This interpretation is consistent with our previous work, which demonstrated that residual stress and biaxial mechanical properties evolve differently during aneurysm progression^12^. These findings suggest that different mechanical descriptors capture distinct aspects of aneurysmal remodeling and therefore provide complementary insight into disease progression. Finally, the substantial unexplained variability observed in the regression models further suggests that aneurysmal mechanics are influenced by additional biological and structural factors not captured by routine clinical characteristics.

## 5 Acknowledgments and Declarations

The authors thank Reza Jafar for his contribution to specimen collection and processing, and Asher Okung, Ayse Hafsa Es-Anderson, and Nihal Yapici for their contributions to the postprocessing of opening-angle and biaxial tensile testing data. The authors are also grateful to the University of Ottawa Cardiac Endowment Fund for supporting this work at the University of Ottawa Heart Institute.

## 6 Declaration of generative AI and AI-assisted technologies

During preparation of this manuscript, the authors used ChatGPT (OpenAI) to assist with language and readability. All generated suggestions were reviewed and edited by the authors, who take full responsibility for the content of the final manuscript.

## References

1. Davies RR, Goldstein LJ, Coady MA, et al. Yearly rupture or dissection rates for thoracic aortic aneurysms: simple prediction based on size. Ann Thorac Surg. 2002;73(1):17–28. doi:10.1016/S0003-4975(01)03236-2

2. Geisbüsch S, Kuehnl A, Salvermoser M, Reutersberg B, Trenner M, Eckstein HH. Editor’s Choice – Hospital Incidence, Treatment, and In Hospital Mortality Following Open and Endovascular Surgery for Thoraco-abdominal Aortic Aneurysms in Germany from 2005 to 2014: Secondary Data Analysis of the Nationwide German DRG Microdata. Eur J Vasc Endovasc Surg. 2019;57(4):488–498. doi:10.1016/j.ejvs.2018.10.030

3. Wu J, Zafar MA, Liu Y, et al. Fate of the unoperated ascending thoracic aortic aneurysm: three-decade experience from the Aortic Institute at Yale University. Eur Heart J. 2023;44(43):4579–4588. doi:10.1093/eurheartj/ehad148

4. Isselbacher EM, Preventza O, Hamilton Black J, et al. 2022 ACC/AHA Guideline for the Diagnosis and Management of Aortic Disease: A Report of the American Heart Association/American College of Cardiology Joint Committee on Clinical Practice Guidelines. Circulation. 2022;146(24):e334-e482. doi:10.1161/CIR.0000000000001106

5. Boczar KE, Cheung K, Boodhwani M, et al. Sex Differences in Thoracic Aortic Aneurysm Growth: Role of Aortic Stiffness. Hypertension. 2019;73(1):190–196. doi:10.1161/HYPERTENSIONAHA.118.11851

6. Chung JC Y., Wong E, Tang M, et al. Biomechanics of Aortic Dissection: A Comparison of Aortas Associated With Bicuspid and Tricuspid Aortic Valves. J Am Heart Assoc Cardiovasc Cerebrovasc Dis. 2020;9(15):e016715. doi:10.1161/JAHA.120.016715

7. Ferrara A, Morganti S, Totaro P, Mazzola A, Auricchio F. Human dilated ascending aorta: Mechanical characterization *via* uniaxial tensile tests. J Mech Behav Biomed Mater. 2016;53:257–271. doi:10.1016/j.jmbbm.2015.08.021

8. Lin S, Morgant MC, Marín-Castrillón DM, et al. Aortic local biomechanical properties in ascending aortic aneurysms. Acta Biomater. 2022;149:40–50. doi:10.1016/j.actbio.2022.06.019

9. Belz GG. Elastic properties and Windkessel function of the human aorta. Cardiovasc Drugs Ther. 1995;9(1):73–83. doi:10.1007/BF00877747

10. Lissoni V, Schembri M, Luraghi G, et al. The Effect Of Localized Circumferential Residual Stress On Pressurized Stress State Of CT Reconstructed Vessels: A Finite Element Study. Ann Biomed Eng. Published online March 23, 2026. doi:10.1007/s10439-026-04085-x

11. Labrosse MR, Gerson ER, Veinot JP, Beller CJ. Mechanical characterization of human aortas from pressurization testing and a paradigm shift for circumferential residual stress. J Mech Behav Biomed Mater. 2013;17(Supplement C):44–55. doi:10.1016/j.jmbbm.2012.08.004

12. Ghadie N, Labrosse M, Boodhwani M. Biomechanical Remodeling of Ascending Aortic Aneurysms Across the 5-cm Surgical Threshold. Submitt Publ. 2026.

13. Mammoto A, Matus K, Mammoto T. Extracellular Matrix in Aging Aorta. Front Cell Dev Biol. 2022;10:822561. doi:10.3389/fcell.2022.822561

14. Subodh V, Siu SC. Aortic Dilatation in Patients with Bicuspid Aortic Valve. N Engl J Med. 2014;370(20):1920–1929. doi:10.1056/NEJMra1207059

15. Olsson C, Thelin S, Ståhle E, Ekbom A, Granath F. Thoracic Aortic Aneurysm and Dissection: Increasing Prevalence and Improved Outcomes Reported in a Nationwide Population-Based Study of More Than 14 000 Cases From 1987 to 2002. Circulation. 2006;114(24):2611–2618. doi:10.1161/CIRCULATIONAHA.106.630400

16. Sokolis DP, Iliopoulos DC. Impaired mechanics and matrix metalloproteinases/inhibitors expression in female ascending thoracic aortic aneurysms. J Mech Behav Biomed Mater. 2014;34:154–164. doi:10.1016/j.jmbbm.2014.02.015

17. Labrosse MR, Jafar R, Ngu J, Boodhwani M. Planar biaxial testing of heart valve cusp replacement biomaterials: Experiments, theory and material constants. Acta Biomater. 2016;45:303–320. doi:10.1016/j.actbio.2016.08.036

18. Humphrey JD. Cardiovascular Solid Mechanics: Cells, Tissues, and Organs. Springer Science & Business Media; 2002.

19. Ghadie NM, Labrosse MR, St-Pierre JP. Glycosaminoglycans modulate compressive stiffness and circumferential residual stress in the porcine thoracic aorta. Acta Biomater. 2023;170:556–566. doi:10.1016/j.actbio.2023.08.061

20. Tarraf SA, Kramer B, Vianna E, et al. Lengthwise regional mechanics of the human aneurysmal ascending thoracic aorta. Acta Biomater. 2023;162:266–277. doi:10.1016/j.actbio.2023.03.023

21. Tjahjadi NS, Kim T, Marway PS, et al. Three-dimensional assessment of ascending aortic stiffness, motion, and growth in ascending thoracic aortic aneurysm. Eur Heart J - Imaging Methods Pract. 2025;3(1):qyae133. doi:10.1093/ehjimp/qyae133

22. Okamoto RJ, Wagenseil JE, DeLong WR, Peterson SJ, Kouchoukos NT, Sundt TM. Mechanical Properties of Dilated Human Ascending Aorta. Ann Biomed Eng. 2002;30(5):624–635. doi:10.1114/1.1484220

23. Sokolis DP. Effects of aneurysm on the directional, regional, and layer distribution of residual strains in ascending thoracic aorta. J Mech Behav Biomed Mater. 2015;46:229–243. doi:10.1016/j.jmbbm.2015.01.024

24. Choudhury N, Bouchot O, Rouleau L, et al. Local mechanical and structural properties of healthy and diseased human ascending aorta tissue. Cardiovasc Pathol. 2009;18(2):83–91. doi:10.1016/j.carpath.2008.01.001

25. Nethononda RM, Lewandowski AJ, Stewart R, et al. Gender specific patterns of age-related decline in aortic stiffness: a cardiovascular magnetic resonance study including normal ranges. J Cardiovasc Magn Reson. 2015;17(1):20. doi:10.1186/s12968-015-0126-0

26. Saini A, Berry C, Greenwald S. Effect of Age and Sex on Residual Stress in the Aorta. J Vasc Res. Published online 1995. doi:10.1159/000159115

